# *CometAnalyser*: a user-friendly, open-source deep-learning microscopy tool for quantitative comet assay analysis

**DOI:** 10.1101/2022.07.04.498642

**Authors:** Attila Beleon, Sara Pignatta, Chiara Arienti, Antonella Carbonaro, Peter Horvath, Giovanni Martinelli, Gastone Castellani, Anna Tesei, Filippo Piccinini

## Abstract

Comet assay provides an easy solution to estimate DNA damage in single cells through microscopy assessment. It is widely used in the analysis of genotoxic damages induced by radiotherapy or chemotherapeutic agents. DNA damage is quantified at the single-cell level by computing the displacement between the genetic material within the nucleus, typically called “comet head”, and the genetic material in the surrounding part of the cell, considered as the “comet tail”. Today, the number of works based on Comet Assay analyses is really impressive. In this work, besides revising the solutions available to obtain reproducible and reliable quantitative data, we developed an easy-to-use tool named *CometAnalyser*. It is designed for the analysis of both fluorescent and silver-stained wide-field microscopy images and allows to automatically segment and classify the comets, besides extracting Tail Moment and several other intensity/morphological features for performing statistical analysis. *CometAnalyser* is an open-source deep-learning tool. It works with Windows, Macintosh, and UNIX-based systems. Source code, standalone versions, user manual, sample images, video tutorial and further documentation are freely available at: *https://sourceforge.net/p/cometanalyser*.

**HIGHLIGHTS:**

1. Comet assay provides an easy solution to estimate DNA damage in single cells.
2. Today, an impressive number of works are based on Comet Assay analyses, especially in the field of cancer research.
3. Comet assay was originally performed as a qualitative analysis.
4. None of the free tools today available work on both fluorescent- and silver-stained images.
5. We developed CometAnalyser, an open-source deep-learning tool designed for easy segmentation and classification of comets in fluorescent- and silver-stained images.

## 1 INTRODUCTION

Comet Assay is a sensitive *in vitro* method to assess DNA damages in individual cells based on the technique of microgel electrophoresis [1]. This assay was originally described by *Ostling* and *Johanson* in 1984 [2]. Its first significant variation, which is the most common today, was proposed 4 years later by *Singh et al*. [3]. The term “Comet” was introduced by *Olive et al*. in 1990 [4] to describe the shape of the DNA visualised upon observing agarose gels. Besides they introduced the concept of “Tail Moment”, computed as the product of two other features, “Tail Length” and “Tail Fluorescence Percentage”, which are some of the most important features considered upon evaluating this assay.

Today, an impressive number of works are based on Comet Assay analyses, especially in the field of cancer research where it is largely used to evaluate different DNA damages induced by ionising radiation or anticancer agents [5]. This significant interest has led to the foundation of an international interest group that offers information, protocols and a forum for discussions on Comet Assay (*https://cometassay.com*, [6]). In a Comet Assay, cells embedded in agarose are: (*a*) lysed, (*b*) subjected to an electric field, (*c*) fluorescent/silver stained, and (*d*) observed using a fluorescence/brightfield microscope. As broken DNA migrates farther in the electric field than the non-damaged genetic material, the cells harbouring DNA damage resemble a “comet” (**Fig. 1a**) with a near-spherical head and a tail region, the latter increasing as DNA damage increases [7].

**Fig. 1.**
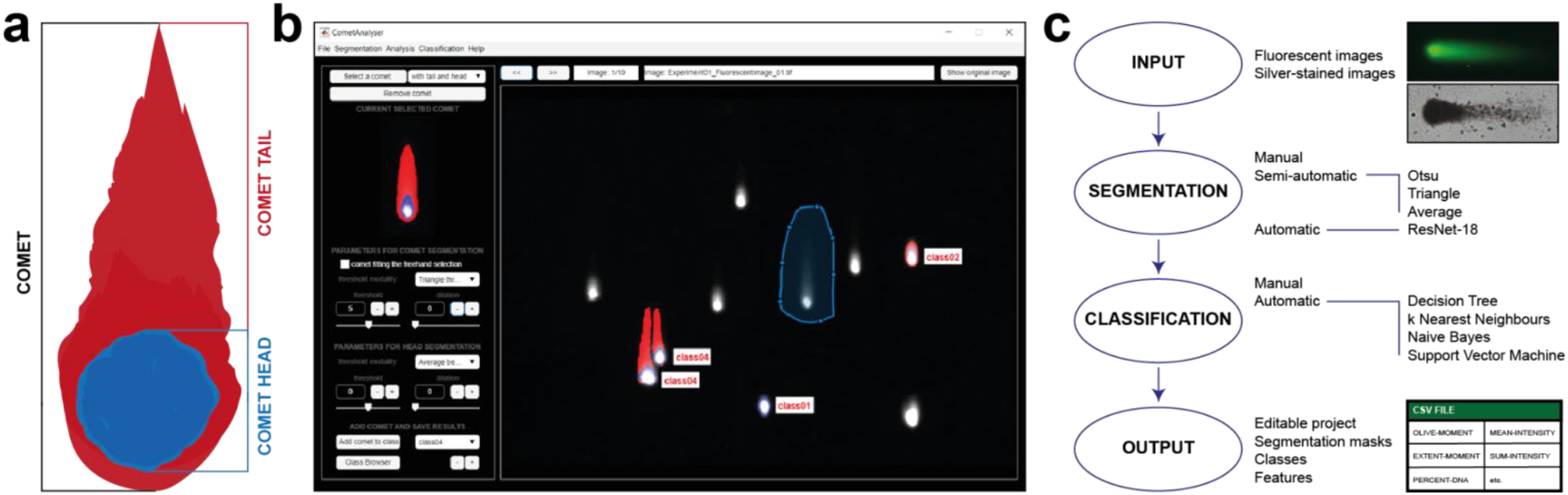
*CometAnalyser*. (*a*) In a Comet Assay, cells resemble comets, with the length of the tails increasing as DNA damage increases. (*b*) *CometAnalyer*’s GUI. (*c*) Flow chart summarising the main possibilities available and steps performed when using CometAnalyser to obtain quantitative results.

Originally, comet assay was performed as a qualitative analysis, but nowadays commercial and freely available tools are available to obtain reproducible quantitative data. However, none of the freely available tools works on both fluorescent- and silver-stained images, which would allow the user to easily segment the comets, and by utilising machine learning methods, automatically identify the most appropriate class for the unannotated comets before extracting several intensity/morphological features, and save the segmentations for future analysis.

In this work, besides revising the existing solutions for performing Comet Assay analysis, we present *CometAnalyser*, an open-source deep-learning tool designed for easy segmentation and classification of comets in fluorescent- and silver-stained images. Source code, standalone versions, user manual, sample images, video tutorial and further documentation are freely available at: *https://sourceforge.net/p/cometanalyser*.

The next sections are organized as follows: **Sect. 2** presents a short overview of the tools designed for performing quantitative analyses of Come Assay microscopy images. **Sect. 3** provides a detailed description of *CometAnalyser*, **Sect. 4** describes the material used and the results obtained in the experiments performed to validate the method proposed. Finally, **Sect. 5** summarises the main findings of the work.

## 2 AVAILABLE TOOLS

In this Section we briefly report a description for *CASP* [8], *CellProfiler* [9] [10], *CometScore* [11], *HiComet* [12], *OpenComet* [13], the tools today freely available for performing quantitative Comet Assay analyses. **Figure 2** reports a representative print-screen of the Graphical User Interface (GUI) for each of them, whilst their main features are summarised in **Table 1** and the links for download in **Table 2**. In addition, **Table 3** reports a reference publication for *CoMat* [14], *CometQ* [15], *DeepComet* [16], *LACAAS* [17], and *SCGE-ProSoftware* [18], tools mentioned in the literature but today not downloadable/available, and **Table 4** the links of the commercial tools today accessible for quantitative Comet Assay analysis, but not freely available for the community.

**Fig. 2.**
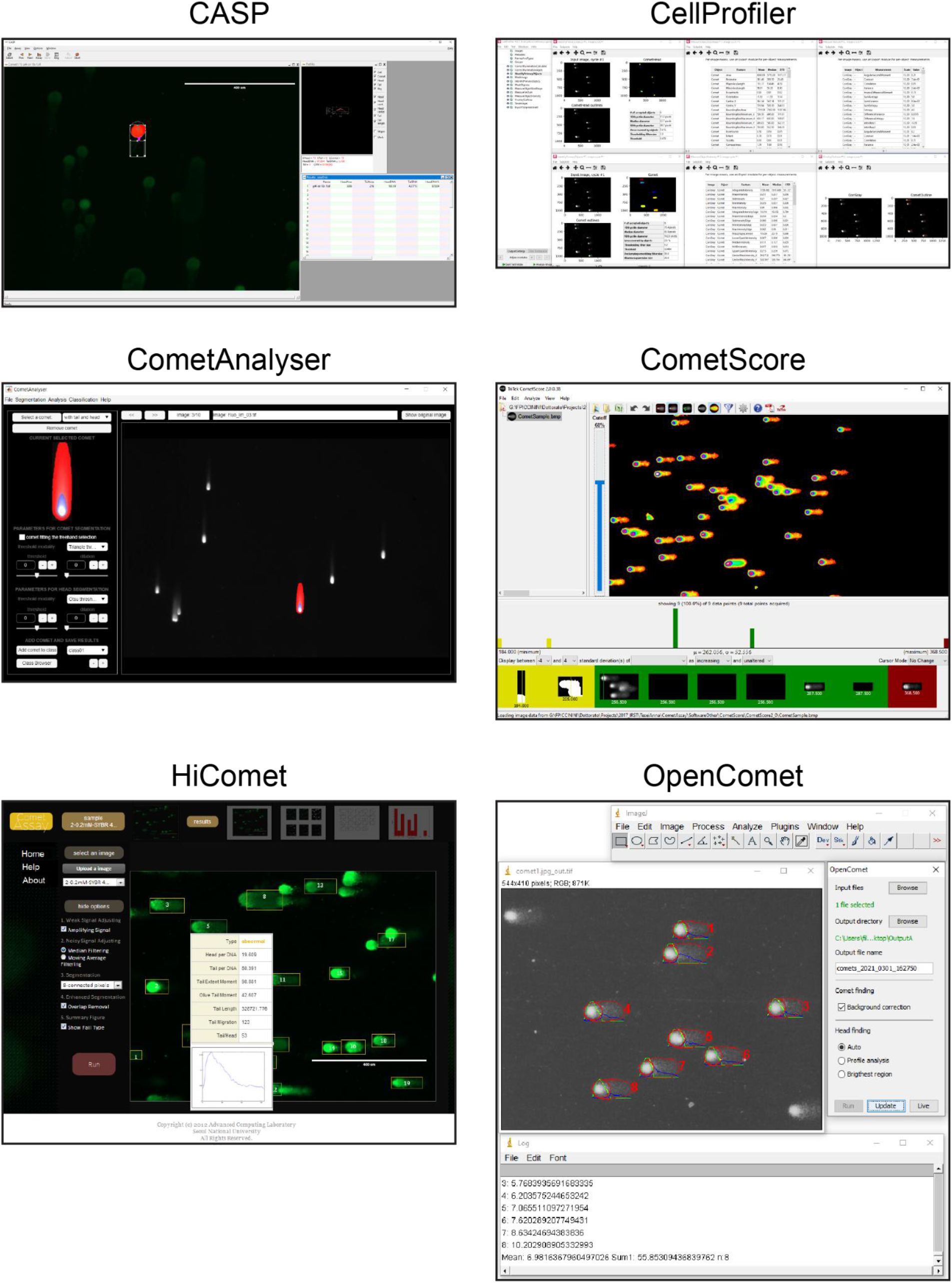
Tools freely available - main window screenshots.

**Table 1:** Tools freely available - characteristics (X = available/yes; O = not available/no).

|  | CASP | CellProfiler | CometAnalyser | CometScore | HiComet | OpenComet |
| --- | --- | --- | --- | --- | --- | --- |
| **VERSION** |  |  |  |  |  |  |
| *Year first release* | 2003 | 2012 | 2021 | 2005 | 2018 | 2014 |
| *Current version* | 1.2.3 beta2 | O | 1.0 | 2.0 | O | 1.3.1 |
| **DOCUMENTATION** |  |  |  |  |  |  |
| *User guide* | X | X | X | X | O | X |
| *Website* | X | X | X | X | O | X |
| *Video tutorial* | O | O | X | X | O | X |
| *Sample dataset* | O | X | X | X | X | O |
| *Open source* | X | X | X | O | X | X |
| *Implementation language* | C++ | Python | Matlab | O | Matlab & PHP | Java |
| **USABILITY** |  |  |  |  |  |  |
| *Input image format* | TIF only | All common | All common | BMP only | All common | All common |
| *No programming experience required* | X | O | X | X | O | X |
| *User-friendly GUI* | X | X | X | X | O | X |
| *Intuitive visualisation settings* | O | X | X | X | O | X |
| *No commercial licences required* | X | X | X | X | X | X |
| *Portability on Win/Linux/Mac* | X | X | X | Win only | X | X |
| **FUNCTIONALITY** |  |  |  |  |  |  |
| *Sylver-stained images* | X | X | X | O | O | O |
| *Fluorescence images* | X | X | X | X | X | X |
| *Automatic comet segmentation* | O | X | X | X | X | X |
| *Semi-automatic comet segmentation* | X | O | X | X | O | O |
| *Manual comet segmentation/correction* | O | O | X | X | O | O |
| *Feature selection* | X | O | X | X | O | O |
| *Automatic comet classification* | O | O | X | X | X | O |
| **OUTPUT** |  |  |  |  |  |  |
| *Segmentation masks* | O | X | X | O | X | O |
| *Feature values* | X | X | X | X | X | X |
| *Classification statistics* | O | O | X | X | X | O |
| *Editable project* | X | O | X | O | O | O |

**Table 2:** Tools freely available - links (last access. 27/06/2022).

|  |  |
| --- | --- |
| **CASP** | http://www.casp.of.pl |
| **CellProfiler (pipeline A)** | https://cellprofiler.org/examples |
| **CellProfiler (pipeline B)** | https://ars.els-cdn.com/content/image/1-s2.0-S1383571812002288-mmc1.pdf |
| **CometAnalyser** | https://sourceforge.net/p/cometanalyser |
| **CometScore** | http://rexhoover.com/index.php?id=cometscore |
| **HiComet** | https://github.com/taehoonlee/hicomet |
| **OpenComet** | http://cometbio.org |

**Table 3.**
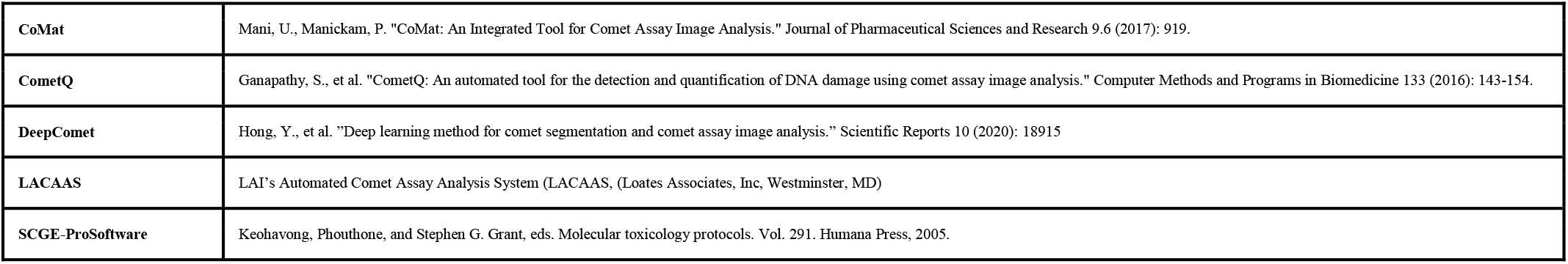
Tools mentioned in the literature but today not downloadable/available - references.

|  |  |
| --- | --- |
| CoMat | Mani, U., Manickam, P. "CoMat: An Integrated Tool for Comet Assay Image Analysis." Journal of Pharmaceutical Sciences and Research 9.6 (2017): 919. |
| CometQ | Ganapathy, S., et al. "CometQ: An automated tool for the detection and quantification of DNA damage using comet assay image analysis." Computer Methods and Programs in Biomedicine 133 (2016): 143-154. |
| DeepComet | Hong, Y., et al. "Deep learning method for comet segmentation and comet assay image analysis." Scientific Reports 10 (2020): 18915 |
| LACAAS | LAI's Automated Comet Assay Analysis System (LACAAS, (Loates Associates, Inc, Westminster, MD) |
| SCGE-ProSoftware | Keohavong, Phouthone, and Stephen G. Grant, eds. Molecular toxicology protocols. Vol. 291. Humana Press, 2005. |

**Table 4.** Commercial tools today available- links (last access 27/06/2022).

|  |  |
| --- | --- |
| Andor - Komet 7.0 | https://andor.oxinst.com/products/komet-software/komet-7 |
| Helix3 | https://www.helix3inc.com |
| Imstar- SCREENTOX | http://www.imstarsa.com/products-applications/ |
| Instem - Comet Assay IV | http://www.scorecomets.com |
| MetaSystems - Metafer | https://metasystems-international.com/en/products/solutions/comet-foci/ |
| Trevigen - Comet Analysis Software | https://trevigen.com/docs/protocol/manual_Trevigen%20Comet%20Analysis%20Software.pdf |

**Table 4.**
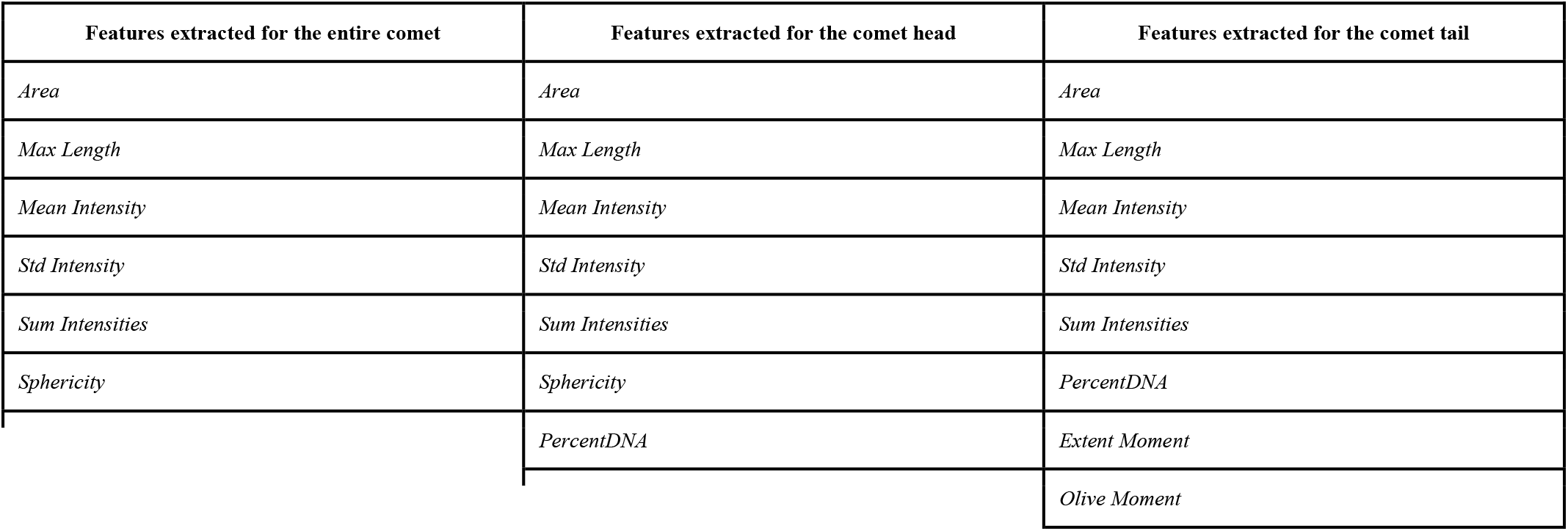
Intensity/morphological features automatically extracted by *CometAnalyser*.

| Features extracted for the entire comet | Features extracted for the comet head | Features extracted for the comet tail |
| --- | --- | --- |
| *Area* | *Area* | *Area* |
| *Max Length* | *Max Length* | *Max Length* |
| *Mean Intensity* | *Mean Intensity* | *Mean Intensity* |
| *Std Intensity* | *Std Intensity* | *Std Intensity* |
| *Sum Intensities* | *Sum Intensities* | *Sum Intensities* |
| *Sphericity* | *Sphericity* | *PercentDNA* |
|  | *PercentDNA* | *Extent Moment* |
|  |  | *Olive Moment* |

### CASP [8]

*CASP* is a user-friendly *C/C++* open-source tool developed by *Końca et al*. in 2003. It works with either silver-stained or fluorescence-stained comets saved in “.tif” format. An unlimited number of images can be marked, and the program will load them successively into an “image view” window. To recognize the comet head and tail it assumes that the most intensive points are placed in the head and the comet is oriented from the left-hand side (head) to the right-hand side (tail) of the image. Accordingly, only comets oriented from left to right can be analysed correctly, but images can be pre-rotated. The tool is a semi-automatic one. Practically, to proceed with the analysis, the user must define a rectangle around the comet of interest. Then, he/she can adjust various thresholds of sensitivity and save the adjustments for future use. An intensity profile of the currently selected comet shows up on a “profile” window together with selected feature values (e.g. tail moment and several other features). When measurements are terminated, the feature values are exported into a text file and the full project can be saved and loaded back for future analysis.

### CellProfiler [9] [10]

*CellProfiler* is a popular *MATLAB*/*Python* software suite worldwide used to set several microscopy image-based analyses by aligning in a pipeline customised modules for standard image processing tasks. Among the different analyses, the comet assay is a typical one, directly reported also in the list of the common applications on the *CellProfiler*’s website. Over the years, several *CellProfiler*’s pipelines have been designed by different groups to optimise the comet assay analysis for different scenarios, including both the silver-stained and fluorescence images. Furthermore, the pipelines can be easily modified with a limited programming experience, and the segmentation masks and the features computed can be easily exported in different formats.

### CometScore [11]

*CometScore* is a freely available software tool developed by *Rex Hoover* in 2005, then supplied in an extended PRO version by the *TriTek Corporation* (Sumerduck, VA). *CometScore* is developed for *Windows* systems only and requires as the input “.bmp” images where all comets are oriented with head on the left and tail on the right. It provides fully-automatic, semi-automatic and manual methods to segment comets based on intensity thresholds defined by the user. Head and tail profiles are visualised directly on the images. *CometScore* also provides an intuitive wizard for comet classification. Feature values can be exported into a text file.

### HiComet [12]

*HiComet* is an automated tool for high-throughput comet-assay analysis. It was developed in *MATLAB* and *PHP* by *Lee et al*. with the idea to freely provide an online implementation to analyse fluorescence comet assay images. It is optimised for rapidly recognising and characterising a large number of comets using little user intervention. It is based on a histogram-thresholding technique for automatically segmenting the comets and no manual correction opportunity is provided to modify the obtained layouts. The feature values appear in a pop-up window shown directly on the image and the comets are then automatically classified on the basis of the features extracted. However, unfortunately, there is no user guide and info about how to run the code available on the *GitHub* repository, no standalone version available, and the Corresponding Author of the reference paper said that today *HiComet* is not maintained anymore. Accordingly, although *HiComet* seems a very promising tool, it is very hard to run it.

### OpenComet [13]

*OpenComet* is a popular *ImageJ*/*Fiji* plugin implemented by *Gyori et al*. in 2014. It is extremely easy-to-use but it works with fluorescence images only where all comets are oriented with head on the left and tail on the right. It requires just the definition of the input and output folders and a few pre-processing decisions. It uses a robust method for finding comets based on geometric shape attributes and segmenting the comet heads through image intensity profile analysis. Head and tail profiles are then visualised directly on the images, while feature values are exported into a spreadsheet file. No manual correction opportunity is provided to modify the comet layout, but after the automated analysis is complete, the user has the option to review the images and click on any comet to remove it from the output if needed. Finally, images with overlaid profiles and the spreadsheet files are saved automatically in the chosen output folder.

## 3 CometAnalyser

*CometAnalyser* is written in *MATLAB*. It provides an easy solution for accurate comet segmentations and quantitative analysis. It exploits advanced processing methods with minimal user interaction to quantitatively evaluate the cell’s viability by assessing DNA damage. This is quantified by computing several intensity/morphological features for every single comet to evaluate the displacement between the genetic material within the nucleus, *i*.*e*. the “comet head”, and the genetic material in the surrounding part, considered as the “comet tail”. An early command-line version of our tool has been used by *Pignatta et al*. for studying DNA damage of cancer cells treated according to different radiotherapy protocols [19]. Since then *CometAnalyser* has been extended by (*I*) implementing machine learning methods to automatically segment and classify the comets; (*II*) developing modules to analyse/modify the segmentations; and (*III*) designing a user-friendly GUI (**Fig. 1b**) subdivided into 4 main modules: (*a*) image/project processing, (*b*) comet segmentation, (*c*) comet classification, (*d*) feature extraction and data sharing (**Fig. 1c**).

### 3.1 Comet segmentation: Fully-automatic modality

Comets are segmented by exploiting a deep convolutional neural network [20]. Pre-trained segmentation networks for both fluorescent- and silver-stained images are provided, and an easy-to-use wizard guides the user in training more specific segmentation models. To create the models we used a built-in *MATLAB* implementation of a convolutional neural network 18 layers deep known in the literature by the name “*ResNet-18*”. Additional details on the network’s architecture are reported at: *https://www.mathworks.com/help/deeplearning/ref/resnet18.html*.

To train the models we used two different datasets of images (**Fig. 3**) with these characteristics: (*a*) *Fluorescent-stained dataset*: The dataset is composed of 33 1200×1600 pixels “.tif” images, containing 20-100 comets for each image. The comets were manually annotated by an expert microscopy user. The slides were prepared according to a standard neutral comet assay manufacturer’s protocol (Comet assay, Trevigen, Gaithersburg, MD). Briefly, 5000 cancer cells (A549, Non Small Cell Lung Cancer cell line, American Type Culture Collection, ATCC, Rockville, MD, USA) were suspended in LMAgarose at 37 °C and immediately transferred onto the comet slide. The slides were immersed for 1 hour at 4 °C in a lysis solution, washed in the dark for 1 hour at room temperature in a neutral solution, and electrophoresed for 30 min at 21 V. Slides were then dipped in 70 % ethanol and stained with the Syber Green (Bio-Rad Laboratories, Hercules, CA, USA). Images were captured using an AMG EVOS FL microscope (Thermo Fisher Scientific, Waltham, MA, USA), equipped with a Sony ICX285AL CCD camera (Tokyo, Japan) at 10× magnification. (*b*) *Silver-stained dataset*: The dataset is composed of 54 1280×1024 “.tif” images, containing 2-20 comets for each image. The comets were manually annotated by an expert microscopy user. The slides were prepared according to a standard alkaline comet assay manufacturer’s protocol (Comet assay, Trevigen, Gaithersburg, MD). Briefly, 5×10^5^ fibroblast cells (normal fibroblast lung cell line MRC-5, American Type Culture Collection, ATCC, Rockville, MD, USA) were suspended in LMAgarose (at 37 °C) and immediately transferred onto the comet slide. The slides were immersed for 1 h at 4 °C in a lysis solution, washed in the dark for 1 h at room temperature in an alkaline solution, and electrophoresed for 30 min at 21 V. Slides were then dipped in 70% ethanol and stained with the Silver Staining Kit (Trevigen) according to the manufacturer’s protocol. Images were captured using an Olympus IX51 microscope (Tokyo, Japan), equipped with a Nikon DS-Vi1 camera (Tokyo, Japan) at 10× magnification.

**Fig. 3.**
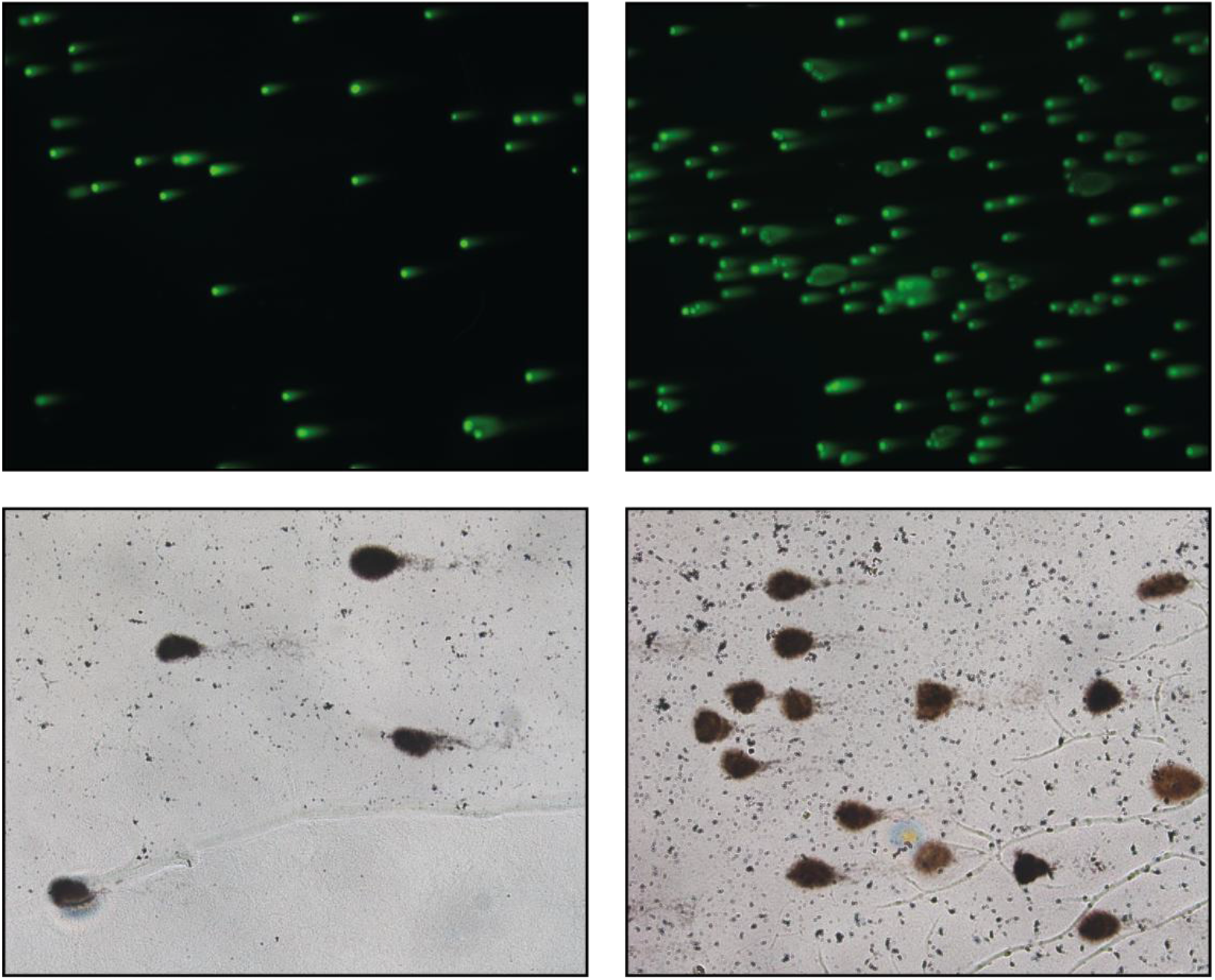
Representative images of the two datasets used for training the deep-learning segmentation models.

The deep learning pre-trained segmentation models created, the images used in the training sets and segmentation masks obtained are freely available at: *https://sourceforge.net/p/cometanalyser*. It is worth noting that the provided pre-trained deep-learning segmentation models logically reliably perform just on images with characteristics similar to those of the images used for training the models. However, the user can easily train a new segmentation model following these steps: (*1*) Annotate an appropriate number of comets. Please note that all the comets in the images included in the training set must be segmented (*i*.*e*. images in the training set cannot be just partially segmented). (*2*) Export the segmentation masks using the *‘‘Segmentation - Export Annotation*’’ button of *CometAnalyser*. (*3*) Train a new deep-learning model using the *‘‘Segmentation - Train New Model’*’ button of *CometAnalyser*. Network parameters can be modified using the *‘‘Segmentation – Training Options’’* button.

### 3.2 Comet segmentation: Semi-automatic modality

Currently, 3 different threshold-based semi-automatic modalities have been implemented in *CometAnalyser*. All of them are based on the analysis of the histogram of the grey-level intensities of the local ROI surrounding the comet to be segmented. The first is the classical *Otsu thresholding* segmentation [21]. The second is the *triangle segmentation* defined by *Zack et al*. [22]. Finally, we implemented an additional segmentation strategy called “*Average between Otsu and Triangle*”, with the final threshold defined as the average value of the ones defined according to the two methods just cited. To segment a comet using one of the semi-automatic modalities available, first the user should simply surround the single comets by drawing a circle. Then, he/she has to select one of the available threshold modalities for segmenting the comet from the background and the head from the tail. Finally, using additional commands for scaling the thresholds and dilating the segmentation masks, he/she can adjust the contours proposed for the head and the comet in general. Once defined the parameters, the settings can be reused for the segmentation of similar comets. *Semi-automatic modality*: the user simply marks single comets by drawing circles.

### 3.3 Comet segmentation: Manual modality

By switching on the “*comet fitting the freehand selection*” check-box present on the left side of the *CometAnalyser*’s GUI, the user enables the manual segmentation modality and can precisely define the border of the comet by directly drawing the corresponding contour.

### 3.4 Comet classification

In the literature, comets are typically classified into 5 different categories, with the first class representing undamaged cells (*i*.*e*. comets with no or barely detectable tails), while class 5 includes just comet tails without visible nuclei [23]. To increase flexibility by providing the opportunity to define sub-classes or perform regression analysis [24], *CometAnalyser* allows the user to define an infinite number of classes. Accordingly, the user can manually define the class for each segmented comet, or may use built-in machine learning algorithms to train a classifier based on previously classified comets, and then use it to automatically define the class of newly segmented ones. It is worth noting that through *CometAnalyser*, the user can easily train a new classifier (*i*.*e*. classification network) following these steps: (*1*) Define the classes. (*2*) Manually classify an appropriate number of comets for each class. (*3*) Export the features (*i*.*e*. “Analysis – Export Measurements” button of *CometAnalyser*) of the classified comets. (*4*) Define the machine learning algorithm to be used for creating the network. Currently, *CometAnalyser* provides four different machine learning algorithms to define the classification network: “*Decision Tree*”, “*k Nearest Neighbours*”, “*Naive Bayes*”, “*Support Vector Machine*”. The machine learning algorithms implemented in *CometAnalyser* are based on *MATLAB* built-in functions. A more detailed description of them is available at: *https://it.mathworks.com/help/stats/classification.html?s_tid=CRUX_lftnav*.

### 3.5 Feature extraction and data sharing

Typically, several intensity/morphological features are evaluated in a Comet Assay analysis [13]. **Table 5** reports the list for the 21 features extracted automatically by *CometAnalyser* separately considering the comet’s head, the comet’s tail, or for the whole comet. Briefly:

**Table 5.** Jaccard Index values (the better value for each row reported in Italics).

| Jaccard Index | CometAnalyser | CellProfiler |
| --- | --- | --- |
| Image 01 | *0.7235* | 0.6611 |
| Image 02 | 0.6598 | *0.6730* |
| Image 03 | *0.6039* | 0.5464 |
| Image 03 | *0.8121* | 0.5690 |
| Image 04 | *0.7280* | 0.3952 |
| Image 05 | *0.7446* | 0.6129 |
| **Image 06** | 0.5773 | *0.6124* |
| **Image 07** | 0.7146 | *0.7982* |
| **Image 08** | 0.5577 | *0.7798* |
| **Image 09** | *0.6723* | 0.6721 |
| **Average Jaccard Index** [mean ± std] | *0.6794 ± 0.1110* | 0.6320 ± 0.1528 |

- *Area* [pixels]: number of pixels belonging to the object.
- *Max Length* [pixels]: length in pixels of the maximum diameters passing through the centroid of the object.
- *Mean Intensity* [grey levels]: average of the intensity values of the pixels of the object.
- *Std Intensity* [grey levels]: standard deviation of the intensity values of the pixels of the object.
- *Sum Intensities* [grey levels]: sum of the intensity values of the pixels of the object.
- *Sphericity* [range 0-1]: sphericity value computed according to the following equation:

~~~
Sphericity = pi * sqrt(a*Area/pi) / Perimeter
~~~
- *PercentDNA* [%]: percentage between *Sum Intensities* of the object (i.e. head or tail) and *Sum Intensities* of the entire comet:

~~~
PercentDNA = 100 * (Object_Sum_Intensities / Comet_Sum_Intensities)
~~~
- *Extent Moment*: ratio between *Sum Intensities* of the comet tail and *Sum Intensities* of the entire comet, multiplied by the *Max Length* of the comet tail:

~~~
Extent_Moment = Tail_Max_Length * (Tail_Sum_Intensities / Comet_Sum_Intensities)
~~~
- *Olive Moment:* ratio between *Sum Intensities* of the comet tail and *Sum Intensities* of the entire comet, multiplied per the absolute value of the module of the distance between the centroid of the head and the centroid of the tail:

~~~
Olive Moment = | Head_centroid_position - Tail_centroid_position | * (Tail_Sum_Intensities / Comet_Sum_Intensities)
~~~

For an extensive description of the features please check also the “*Appendix: Equations*” in the tutorial of the *CometScore* tool, available at: *https://dokument.pub/ql/cometscore-tutorial-automatic-comet-assay-flipbook-pdf*. Moreover, *CometAnalyser* provides an easy solution for the users to define new features. The extracted features are saved in a spreadsheet file, and the segmentation masks are saved as “.png” files. All the segmented comets are shown as thumbnails in a pop-up wizard called “*Comet Browser*”, and the user can navigate the original images with just a click to find and eventually modify the segmented comets which are highlighted in red (DNA) and blue (nucleus). The project can be saved and shared, allowing future modifications and analysis.

## 4 EXPERIMENTAL RESULTS

*CometAnalyser* was compared with the competitor tools. First, we discarded from the comparison the tools reported in **Table 1** but not providing output masks of the segmented comets (*i*.*e. CASP, CometScore* and *OpenComet*). In addition, we also discarded *HiComet* because today it is not maintained and currently there are no available instructions on how to run the tool. Concluding, we defined a testbed dataset, and using the ground truth, we computed the Jaccard Index of the masks automatically obtained by *CometAnalyser* and *CellProfiler*.

In particular, we analysed the masks of the comets automatically segmented using the fluorescence pre-trained deep-learning segmentation network provided at: *https://sourceforge.net/p/cometanalyser*. First, we defined a dataset of 10 new fluorescent images with characteristics similar to those used for training the segmentation network. Then, we asked an expert microscopist to manually segment the comets for defining the ground truth masks. To evaluate the segmentation quality of the comets’ masks, we computed the Jaccard Index [25]. The values obtained for each image are reported in **Table 5**. Both tools obtained very good segmentation masks (**Fig. 4**) and *CometAnalyser* achieved a slightly better Jaccard Index (*i*.*e*. 0.68 versus 0.63).

**Fig. 4.**
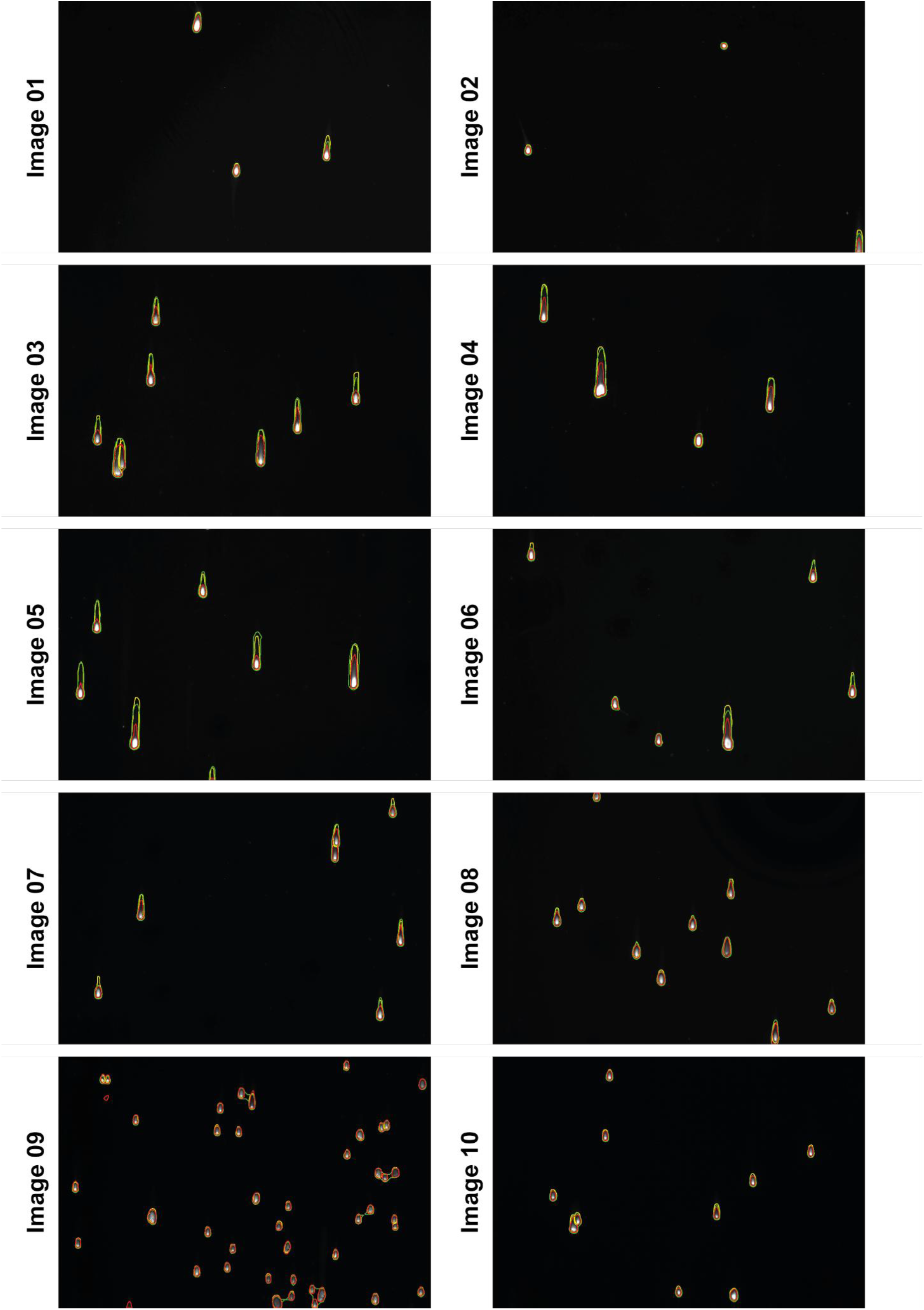
Segmentations. Ground truth (yellow outline), *CometAnalyser*’s masks (green outline), *CellProfiler*’s masks (red outline).

Testbed dataset, ground truth masks, *CometAnalyser*’s masks, *CellProfiler*’s pipeline and masks and the *MATLAB* code to compute the Jaccard Index are provided in a “.zip” folder named: “*CometAnalyser VS CellProfiler*”, available at: *https://sourceforge.net/p/cometanalyser*.

## 5 CONCLUSIONS

*CometAnalyser* is an open-source tool to quantitatively assess DNA damage, for instance to analyse genotoxic effects induced by radio- and chemotherapy. *CometAnalyser* is developed in *MATLAB* and it works for both fluorescence and silver-stained comet assay microscopy images. It provides fully-automatic, semi-automatic and manual methods to segment comets in several scenarios. The main aim of *CometAnalyser* is to perform accurate comet segmentations and quantitative analysis using advanced image processing methods with minimal user interaction. It is well documented with a user manual, a video tutorial, sample images and results a very easy-to-use tool. A deep-learning model for automatic comet segmentation is provided. Furthermore, an intuitive GUI helps the user in defining interesting classes and annotating representative comets. Finally, the projects can be saved and loaded back for future analysis. Concluding, *CometAnalyser* is currently the most complete freely available solution for 3 main reasons: (*a*) it works with both fluorescent- and silver-stained images; (*b*) it provides easy-to-use machine learning modules for the segmentation and classification of the comets; and (*c*) it offers several opportunities to edit the segmentations and export the data.

## ACKNOWLEDGEMENTS

The authors thank Roberto Vespignani and Nicola Caroli (IRST, Meldola, Italy) for technical support; Dora Bokor PharmD (Szeged, Hungary) for the English revision.

## Author Contributions

Conceptualization: AB, SP, CA, AT, FP;

Data curation: SP, CA;

Formal analysis: AB, FP;

Funding acquisition: AC, PH, GM, GC, AT;

Investigation: AB, SP, CA, AT, FP;

Methodology: AB, SP, CA, FP;

Project administration: FP;

Resources: AC, PH, GM, GC, AT;

Software: AB, PH, GC, FP;

Supervision: AC, PH, GM, GC, AT;

Validation: AB, SP, CA, AC, FP;

Visualization: SP, CA, AC;

Writing - original draft: AB, SP, CA, FP;

Writing - review & editing: AC, PH, GM, GC, AT.

## Funding

A.B. and P.H. acknowledge support from the LENDULET-BIOMAG Grant (2018-342), from OTKA-SNN, from TKP2021-EGA09, from H2020-COMPASS-ERAPerMed, from CZI Deep Visual Proteomics, from H2020-DiscovAir, H2020-Fair-CHARM, from the ELKH-Excellence grant, from the Finnish TEKES FiDiPro Fellow Grant 40294/13 and FIMM High Content Imaging and Analysis Unit (FIMMHCA; HiLIFE-HELMI) and Biocenter Finland, Finnish Cancer Society, Juselius Foundation, Academy of Finland Centre of Excellence in Translational Cancer Biology, Kymenlaakso and Finnish Cultural Foundation. F.P. acknowledges support from the Union for International Cancer Control (UICC) for a UICC Yamagiwa-Yoshida (YY) Memorial International Cancer Study Grant (ref: UICC-YY/678329). S.P., C.A., G.M., A.T., and F.P. acknowledges support by the Italian Ministry of Health, contribution “*Ricerca Corrente*” within the research line “*Appropriateness, outcomes, drug value and organizational models for the continuity of diagnostic-therapeutic pathways in oncology*”.

## Conflict of interest statements

The author declares that he has no conflict of interest.

## Data Availability Statement

The data presented in this study are available at: *https://sourceforge.net/p/cometanalyser*

## Notes

https://sourceforge.net/p/cometanalyser

### Competing Interest Statement

The authors have declared no competing interest.

